## Supplementary Information for "Quantifying how post-transcriptional noise and gene copy number variation bias transcriptional parameter inference from mRNA distributions"

**7 Classification of the cell cycle phase: bimodal Gaussian distribution method versus the Fried/Baisch model 20**

### 1 Accuracy of inference from synthetic mature and nascent mRNA data

#### 1.1 Inference from synthetic mature mRNA data with external noise

In the main text, Fig 2, we showed how the addition of 5% external noise to synthetic mature mRNA data degrades the inference accuracy. In Fig. 1 we show how the addition of a larger amount of external noise (10%) causes an even larger loss of accuracy. In particular for 91% of the parameters, the inference accuracy is higher when using nascent mRNA data (Fig. 1a) and the median relative errors become very high for most parameters, especially for $\rho$ and $\sigma_{\text{off}}$ in the limit of large $f_{\text{ON}}$ (Fig. 1b).

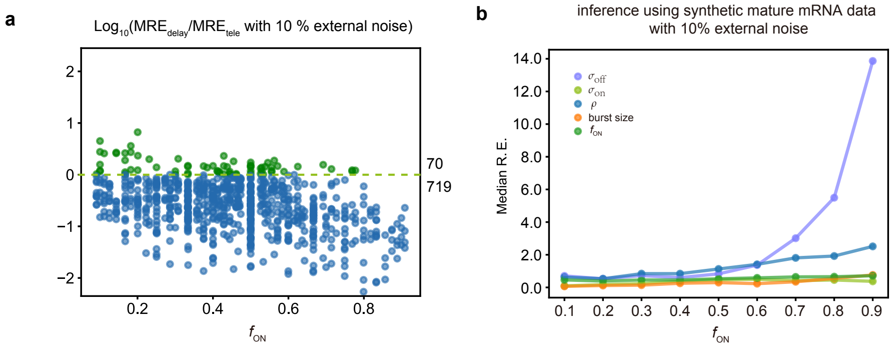

**Figure 1:** Comparing inference accuracy using synthetic nascent mRNA data and synthetic mature mRNA data with 10% external noise (lognormal distributed noise is added to the initiation rate $\rho$ to mimic external noise due to post-transcriptional processing that is only present in mature mRNA). **a.** Ratio of the mean relative errors in the two types of data as a function of the true fraction of ON time, $f_{\text{ON}}$. For $\approx$ 91% (719/789) of the parameters, the inference accuracy is higher when using nascent mRNA data. **b.** The median relative error of each transcriptional parameter as a function of the fraction of ON time using synthetic mature mRNA.

#### 1.2 Accuracy of distribution fits

**a**

| | True | | | | | Telegraph | | | | |
|---|---|---|---|---|---|---|---|---|---|---|
| Parameter | $\sigma_{\text{off}}$ | $\sigma_{\text{on}}$ | $\rho$ | burst size | $f_{\text{ON}}$ | $\sigma_{\text{off}}$ | $\sigma_{\text{on}}$ | $\rho$ | burst size | $f_{\text{ON}}$ |
| Set 1 | 120.38 | 7.70 | 40.85 | 0.34 | 0.06 | 15.58 | 5.63 | 9.22 | 0.59 | 0.27 |
| Set 2 | 35.48 | 17.39 | 85.05 | 2.40 | 0.33 | 36.73 | 18.03 | 85.17 | 2.32 | 0.33 |
| Set 3 | 0.52 | 0.42 | 46.21 | 88.36 | 0.44 | 0.57 | 0.42 | 46.62 | 82.47 | 0.43 |
| Set 4 | 0.32 | 0.47 | 15.71 | 48.65 | 0.59 | 0.33 | 0.47 | 15.70 | 48.01 | 0.59 |
| Set 5 | 0.53 | 5.85 | 49.95 | 94.72 | 0.92 | 0.64 | 6.62 | 50.20 | 77.99 | 0.91 |
| Set 6 | 0.16 | 3.89 | 26.45 | 169.81 | 0.96 | 0.11 | 2.94 | 26.30 | 238.76 | 0.96 |

**b**

| | True | | | | | Delay | | | | |
|---|---|---|---|---|---|---|---|---|---|---|
| Parameter | $\sigma_{\text{off}}$ | $\sigma_{\text{on}}$ | $\rho$ | burst size | $f_{\text{ON}}$ | $\sigma_{\text{off}}$ | $\sigma_{\text{on}}$ | $\rho$ | burst size | $f_{\text{ON}}$ |
| Set 1 | 34.49 | 2.80 | 16.65 | 0.48 | 0.08 | 62.74 | 3.10 | 26.38 | 0.42 | 0.05 |
| Set 2 | 135.68 | 32.98 | 179.74 | 1.32 | 0.20 | 80.42 | 29.44 | 131.35 | 1.63 | 0.27 |
| Set 3 | 9.86 | 7.96 | 63.16 | 6.41 | 0.45 | 7.85 | 6.78 | 61.20 | 7.79 | 0.46 |
| Set 4 | 2.26 | 2.14 | 18.64 | 8.26 | 0.49 | 1.77 | 1.76 | 18.28 | 10.36 | 0.50 |
| Set 5 | 2.27 | 5.80 | 12.55 | 5.53 | 0.72 | 1.60 | 4.48 | 12.21 | 7.63 | 0.74 |
| Set 6 | 1.18 | 9.94 | 24.92 | 21.15 | 0.89 | 0.94 | 8.74 | 24.74 | 26.37 | 0.90 |

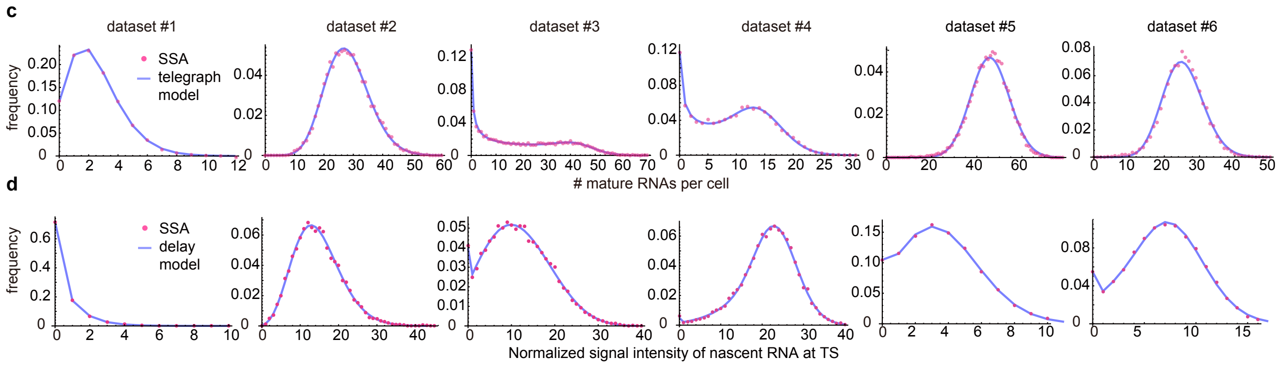

**Figure 2:** **a.** Estimates using the inference algorithm with the telegraph model (with no external noise) for six parameter sets. For both the ground truth and the estimated parameters, we fix the degradation rate $d = 1\ \text{min}^{-1}$. **b.** Estimates using the inference algorithm with the delay telegraph model for six parameter sets. For both the ground truth and the estimated parameters, we fix the delay $\tau = 0.5$ min. **c.** Distributions from synthetic mature mRNA data fitted using the telegraph model. **d.** Distributions from synthetic nascent mRNA data fitted using the delay telegraph model.

In the main text, Fig 2, we showed how the accuracy of parameter estimation is not uniform across parameter space. Here we investigate if there is any relationship between this accuracy and how well is a distribution of mature mRNA numbers / signal intensity fit by the inference algorithm. For 12 parameter sets (6 for the telegraph model – Fig. 2a) and (6 for the delay telegraph model – Fig. 2b), we evaluate the fits to the distribution of synthetic data in Fig. 2 c,d. The results show that independent of the accuracy of the parameters estimated by the inference algorithm, the fits of the delay telegraph and telegraph model distributions to the distributions generated from synthetic data are generally excellent.

#### 1.3 Testing the variability of the inference procedure

To obtain a better understanding of the variability of the inference procedure, for each of the 6 parameter sets in Fig. 2b, i.e. for the inference using the delay telegraph model, we generated 10 independent sets of synthetic data and used maximum likelihood to infer the parameters for each dataset. The mean and standard deviation of the parameters (computed over all 10 datasets) are shown in Table 1. The means are close to the true parameter values (in SI Fig. 2b); this shows that the inference procedure is working correctly and the deviations from ground truth values are mostly due to noise in the synthetic datasets. We can define the sample variability of a parameter as the standard deviation divided by the mean. Ordering parameters by this quantity, we find that for all 6 parameter sets the error is largest for $\sigma_{\text{off}}$. For small $f_{\text{ON}}$, the parameters ordered by increasing error are: $\sigma_{\text{on}}$, burst size, $\rho$, $f_{\text{ON}}$ and $\sigma_{\text{off}}$. While for large $f_{\text{ON}}$, the order is: $\rho$, $f_{\text{ON}}$, $\sigma_{\text{on}}$, burst size and $\sigma_{\text{off}}$. Note that this order is the same as we determined using the relative errors (from the ground truth) which is shown in Fig. 2 of the main text.

| | Mean | | | | | Standard deviation | | | | |
|---|---|---|---|---|---|---|---|---|---|---|
| Parameter | $\sigma_{\text{off}}$ | $\sigma_{\text{on}}$ | $\rho$ | burst size | $f_{\text{ON}}$ | $\sigma_{\text{off}}$ | $\sigma_{\text{on}}$ | $\rho$ | burst size | $f_{\text{ON}}$ |
| Set 1 | 31.85 | 2.69 | 15.68 | 0.53 | 0.10 | 19.07 | 0.22 | 7.68 | 0.07 | 0.05 |
| Set 2 | 92.67 | 30.47 | 141.42 | 1.56 | 0.25 | 23.76 | 1.51 | 21.61 | 0.14 | 0.04 |
| Set 3 | 9.94 | 8.01 | 63.35 | 6.39 | 0.45 | 0.72 | 0.19 | 1.94 | 0.25 | 0.01 |
| Set 4 | 2.30 | 2.16 | 18.76 | 8.17 | 0.48 | 0.12 | 0.05 | 0.31 | 0.31 | 0.01 |
| Set 5 | 2.26 | 5.80 | 12.57 | 5.59 | 0.72 | 0.19 | 0.30 | 0.11 | 0.41 | 0.01 |
| Set 6 | 1.22 | 10.01 | 24.97 | 20.53 | 0.89 | 0.10 | 0.43 | 0.16 | 1.65 | 0.01 |

**Table 1:** Mean and standard deviation of the parameters estimated from 10 independent synthetic datasets, generated for each parameter set in Fig. 2b.

#### 1.4 Confidence intervals using profile likelihood

We perform a profile likelihood study [1] on the 12 parameter sets of synthetic mature/nascent mRNA data described in Section 1.2. We obtain the 95% confidence interval for each parameter. The results are shown in Table 2. We also compare the relative errors for each parameter (computed as (estimated value - ground truth)/ground truth using the data in Fig. 2a and b) with the profile likelihood error (computed as (upper bound - lower bound)/optimal estimate using the bounds in Table 2 and the optimal values in Fig. 2a and b). The results are shown in Table 3. Note that in most cases, the parameters ordered by relative error are in agreement with the parameters ordered by profile likelihood error.

| | Telegraph CI | | | Delay CI | | |
|---|---|---|---|---|---|---|
| Parameter | $\sigma_{\text{off}}$ | $\sigma_{\text{on}}$ | $\rho$ | $\sigma_{\text{off}}$ | $\sigma_{\text{on}}$ | $\rho$ |
| Set 1 | (6.76, 300.00) | (3.53, 8.49) | (5.59, 107.67) | (13.80, 300.00) | (1.85, 2.69) | (9.94, 160.51) |
| Set 2 | (17.22, 190.38) | (14.56, 23.92) | (61.14, 250.51) | (103.80, 268.35) | (30.00, 35.38) | (161.54, 300.00) |
| Set 3 | (0.54, 0.59) | (0.41, 0.43) | (46.08, 47.15) | (6.83, 8.55) | (6.42, 7.04) | (58.28, 63.18) |
| Set 4 | (0.31, 0.35) | (0.46, 0.49) | (15.50, 15.91) | (1.64,1.99) | (1.65,1.85) | (17.92,18.77) |
| Set 5 | (0.49, 0.85) | (5.82, 7.62) | (49.62, 50.92) | (1.56,2.59) | (4.58,5.85) | (12.08,12.97) |
| Set 6 | (0.08, 0.16) | (2.46, 3.54) | (26.10, 26.54) | (0.73,1.24) | (7.53,9.75) | (24.32,25.14) |

**Table 2:** 95% confidence intervals of the 12 parameter sets (shown in Fig. 2a and b).

| | Telegraph | | | | | | Delay | | | | | |
|---|---|---|---|---|---|---|---|---|---|---|---|---|
| | Relative error | | | Profile likelihood error | | | Relative error | | | Profile likelihood error | | |
| | $\sigma_{\text{off}}$ | $\sigma_{\text{on}}$ | $\rho$ | $\sigma_{\text{off}}$ | $\sigma_{\text{on}}$ | $\rho$ | $\sigma_{\text{off}}$ | $\sigma_{\text{on}}$ | $\rho$ | $\sigma_{\text{off}}$ | $\sigma_{\text{on}}$ | $\rho$ |
| Set 1 | 0.87 | 0.27 | 0.77 | 18.82 | 0.88 | 11.07 | 0.82 | 0.11 | 0.58 | 1.75 | 0.34 | 1.82 |
| Set 2 | 0.04 | 0.04 | 1.44E-03 | 4.72 | 0.52 | 2.22 | 0.41 | 0.11 | 0.27 | 0.81 | 0.16 | 0.55 |
| Set 3 | 0.08 | 0.01 | 0.01 | 0.09 | 0.06 | 0.02 | 0.20 | 0.15 | 0.03 | 0.22 | 0.09 | 0.08 |
| Set 4 | 0.01 | 0.01 | 1.11E-03 | 0.14 | 0.07 | 0.03 | 0.22 | 0.18 | 0.02 | 0.20 | 0.11 | 0.05 |
| Set 5 | 0.22 | 0.13 | 4.97E-03 | 0.55 | 0.27 | 0.03 | 0.30 | 0.23 | 0.03 | 0.64 | 0.28 | 0.07 |
| Set 6 | 0.29 | 0.24 | 0.01 | 0.74 | 0.37 | 0.02 | 0.20 | 0.12 | 0.01 | 0.55 | 0.25 | 0.03 |

**Table 3:** Table showing the relative error against profile likelihood error of 12 parameter sets (shown in Fig. 2a and b). See text for details.

#### 1.5 Effect of random perturbation of mature mRNA data on inference

To assess the reliability of the inference results due to errors in spot counting, we redid the inference with synthetic mRNA data (and using the telegraph model) perturbed randomly by minus 1/plus 1/unchanged with probability 1/3. The results are shown in Table 4.

We found that when the fraction of ON time was very small (Set 1), there is a considerable effect of the perturbations on the values of the inferred parameters – this is because in this case, the mean number of mRNA is very small and hence a perturbation of one molecule is very significant. However as expected, the inference results are quite robust when the fraction of ON time is not too small (Sets 2-6).

| | True | | | | | Unperturbed | | | | | -1/0/+1 stochastic perturbation | | | | |
|---|---|---|---|---|---|---|---|---|---|---|---|---|---|---|---|
| Parameter | $\sigma_{\text{off}}$ | $\sigma_{\text{on}}$ | $\rho$ | burst size | $f_{\text{ON}}$ | $\sigma_{\text{off}}$ | $\sigma_{\text{on}}$ | $\rho$ | burst size | $f_{\text{ON}}$ | $\sigma_{\text{off}}$ | $\sigma_{\text{on}}$ | $\rho$ | burst size | $f_{\text{ON}}$ |
| Set 1 | 120.38 | 7.70 | 40.85 | 0.34 | 0.06 | 15.58 | 5.63 | 9.22 | 0.59 | 0.27 | 0.59 | 1.17 | 3.70 | 6.28 | 0.66 |
| Set 2 | 35.48 | 17.39 | 85.05 | 2.40 | 0.33 | 36.73 | 18.03 | 85.17 | 2.32 | 0.33 | 24.13 | 15.79 | 70.89 | 2.94 | 0.40 |
| Set 3 | 0.52 | 0.42 | 46.21 | 88.36 | 0.44 | 0.57 | 0.42 | 46.62 | 82.47 | 0.43 | 0.61 | 0.46 | 47.17 | 76.74 | 0.43 |
| Set 4 | 0.32 | 0.47 | 15.71 | 48.65 | 0.59 | 0.33 | 0.47 | 15.70 | 48.01 | 0.59 | 0.39 | 0.54 | 16.09 | 41.17 | 0.58 |
| Set 5 | 0.53 | 5.85 | 49.95 | 94.72 | 0.92 | 0.64 | 6.62 | 50.20 | 77.99 | 0.91 | 0.68 | 6.72 | 50.35 | 74.48 | 0.91 |
| Set 6 | 0.16 | 3.89 | 26.45 | 169.81 | 0.96 | 0.11 | 2.94 | 26.30 | 238.76 | 0.96 | 0.13 | 3.06 | 26.42 | 203.20 | 0.96 |

**Table 4:** Effects of random perturbations on inference of parameters from mature mRNA data (using the telegraph model).

#### 1.6 Effect of lognormal noise in nascent fluorescent signal on inference

Given the synthetic nascent mRNA signal data $\{S_i\}_{i=1}^{N}$ (where $N$ represents the number of samples) generated by delay telegraph model, for each $S_i$, we perform a random perturbation $\Phi$ under the following conditions

$$\Phi : S_i \to \Phi(S_i)$$

where $\Phi$ is a stochastic perturbation satisfying the following constraints

$$\Phi(S_i) \text{ is a random variable sampled from the distribution LogNormal}(\alpha, \beta)$$
$$\text{whose mean equals } S_i, \text{ and the standard deviation equals } 0.1 * S_i.$$

This means the random perturbation keeps the mean value of the signal $S_i$ unchanged but adds noise with a coefficient of variation equal to 0.1.

In Table 5 we compare the results of inference using synthetic nascent mRNA data with the aforementioned stochastic perturbation and without. As for mature mRNA data, we find that the perturbation has only a significant impact when the fraction of ON time is very small.

| | True | | | | | Delay | | | | | Perturbation | | | | |
|---|---|---|---|---|---|---|---|---|---|---|---|---|---|---|---|
| Parameter | $\sigma_{\text{off}}$ | $\sigma_{\text{on}}$ | $\rho$ | burst size | $f_{\text{ON}}$ | $\sigma_{\text{off}}$ | $\sigma_{\text{on}}$ | $\rho$ | burst size | $f_{\text{ON}}$ | $\sigma_{\text{off}}$ | $\sigma_{\text{on}}$ | $\rho$ | burst size | $f_{\text{ON}}$ |
| Set 1 | 34.49 | 2.80 | 16.65 | 0.48 | 0.08 | 62.74 | 3.10 | 26.38 | 0.42 | 0.05 | 50.08 | 1.52 | 31.71 | 0.63 | 0.03 |
| Set 2 | 135.68 | 32.98 | 179.74 | 1.32 | 0.20 | 80.42 | 29.44 | 131.35 | 1.63 | 0.27 | 233.96 | 30.74 | 300.00 | 1.28 | 0.12 |
| Set 3 | 9.86 | 7.96 | 63.16 | 6.41 | 0.45 | 7.85 | 6.78 | 61.20 | 7.79 | 0.46 | 8.85 | 6.63 | 65.49 | 7.40 | 0.43 |
| Set 4 | 2.26 | 2.14 | 18.64 | 8.26 | 0.49 | 1.77 | 1.76 | 18.28 | 10.36 | 0.50 | 1.90 | 1.64 | 18.65 | 9.83 | 0.46 |
| Set 5 | 2.27 | 5.80 | 12.55 | 5.53 | 0.72 | 1.60 | 4.48 | 12.21 | 7.63 | 0.74 | 2.59 | 4.79 | 13.27 | 5.13 | 0.65 |
| Set 6 | 1.18 | 9.94 | 24.92 | 21.15 | 0.89 | 0.94 | 8.74 | 24.74 | 26.37 | 0.90 | 1.84 | 10.15 | 26.10 | 14.18 | 0.85 |

**Table 5:** Inference using the delay telegraph model from synthetic nascent fluorescent data, with and without perturbation by lognormal noise.

### 2 Telegraph model versus delay telegraph model

In this section, we aim to precisely understand the differences between the telegraph and the delay telegraph models. For both models, we define the rate of switching from the ON state to OFF state as $\sigma_{\text{off}}$, the rate of switching from the OFF state to the ON state as $\sigma_{\text{on}}$ and the production rate of nascent mRNAs in the ON state as $\rho$. The first-order decay rate of nascent mRNA in the telegraph model is given by $d$ and the delay time between initiation and degradation in the delay telegraph model is $\tau$. Note that while the telegraph model was in the main text explained in terms of mature mRNA, in this section we use it as a model for nascent mRNA since we want to compare directly with the delay telegraph model. The telegraph and delay telegraph models can be solved exactly in steady-state conditions [2,3] (an alternative derivation for the delay telegraph model is also given in Section 6). From the generating function solution of the models, one can deduce expressions for the first and second centered moments in steady-state conditions:

$$\begin{aligned}
\langle n\rangle_{\text{tele}} &= \frac{\rho\sigma_{\text{on}}}{d(\sigma_{\text{on}}+\sigma_{\text{off}})},\\
\langle n\rangle_{\text{delay}} &= \frac{\rho\sigma_{\text{on}}\tau}{\sigma_{\text{on}}+\sigma_{\text{off}}},\\
\text{Var}_{\text{delay}} &= \frac{\rho\sigma_{\text{on}}\left((\sigma_{\text{off}}+\sigma_{\text{on}})^3\tau+2\rho\sigma_{\text{off}}\left(e^{-(\sigma_{\text{off}}+\sigma_{\text{on}})\tau}+(\sigma_{\text{off}}+\sigma_{\text{on}})\tau-1\right)\right)}{(\sigma_{\text{off}}+\sigma_{\text{on}})^4},\\
\text{Var}_{\text{tele}} &= \frac{\rho\sigma_{\text{on}}\left(\rho\sigma_{\text{off}}+d(\sigma_{\text{off}}+\sigma_{\text{on}})+(\sigma_{\text{off}}+\sigma_{\text{on}})^2\right)}{d(\sigma_{\text{off}}+\sigma_{\text{on}}+d)(\sigma_{\text{off}}+\sigma_{\text{on}})^2},
\end{aligned} \tag{2.1}$$

where $\langle n\rangle_{\text{delay}}$ and $\langle n\rangle_{\text{tele}}$ are the mean number of nascent mRNA in the delay telegraph and telegraph models respectively, and $\text{Var}_{\text{delay}}$ and $\text{Var}_{\text{tele}}$ are the corresponding variances in molecule numbers. To understand the differences between the two models, we set the means of the two models to be the same (by choosing $d = 1/\tau$) and then compute the relative error in their variance predictions:

$$R = \frac{\text{Var}_{\text{delay}}-\text{Var}_{\text{tele}}}{\text{Var}_{\text{delay}}} = \frac{\left(2(1+x)+e^x(x^2-2)\right)y}{(1+x)(2y+e^x(x+2(x-1)y))}, \tag{2.2}$$

where $x$ and $y$ are non-dimensional variables defined as $x=\tau(\sigma_{\text{off}}+\sigma_{\text{on}})$ and $y=(\rho/\sigma_{\text{off}})(\sigma_{\text{off}}/(\sigma_{\text{off}}+\sigma_{\text{on}}))^2$. Note that $x$ is the ratio of the time for nascent mRNA to detach from the gene $\tau$ and the timescale of promoter switching $1/(\sigma_{\text{off}}+\sigma_{\text{on}})$. The non-dimensional parameter $y$ is a measure of the burstiness of gene expression since it increases with the mean burst size $\rho/\sigma_{\text{off}}$ and the fraction of time spent in the OFF state $\sigma_{\text{off}}/(\sigma_{\text{off}}+\sigma_{\text{on}})$. In fact as $\sigma_{\text{off}}\to 0$ which implies $y\to 0$, the telegraph model converges to a constitutive model where the time between two successive nascent mRNA production events is exponentially distributed. Note that the relative errors in the Fano factor and the coefficient of variation squared are the same as the error in the variance since the means of the two models are the same.

Using Eq. (2.2), it is easy to see that the relative error between the two models vanishes in the limit of $x\to 0$ (when the promoter switching timescales are much longer than the time spent by a polymerase on a gene) or in the limit of $y\to 0$ (when there is no burstiness in gene expression). Hence, the telegraph model is an accurate approximation of the delay telegraph model in these two limits. It can be shown that in the first case, the distribution of nascent mRNA numbers is well approximated by the sum of two Poissons, whereas for the second case the distribution is a Poisson. Note that since $R$ is always positive, it follows that the *telegraph model systematically underestimates the size of noise in nascent mRNA numbers*. It can be further shown that $R$ increases monotonically with $x$ and $y$ and the maximum attainable value is $R=1/2$ (when $x\to\infty, y\to\infty$), i.e., the worst prediction of the telegraph model is that the variance is half that of the delay telegraph model.

In Fig. 3a and Fig. 3c we contrast the distributions of nascent mRNA predicted by the delay telegraph model with those predicted by the telegraph model for the case $d=1/\tau$ where the means are matched, as also assumed for the calculation of the relative error above. Fig. 3b shows the effect of increasing $x$ (via $\tau$) and Fig. 3d shows the effect of increasing $y$ (via $\rho$). The number distributions are constructed from the generating functions of the telegraph model [2] and of the delay telegraph model (Section 6). Note that the shapes of the distributions of the two models can be considerably different, e.g. the cases $\rho = 3$ and $\rho = 10$ in Fig. 3c shows that the nascent distribution from the delay model is bimodal with peaks at 0 and at a non-zero value but it is unimodal with a non-zero peak from the telegraph model. Hence we conclude that if we are interested in accurately predicting nascent mRNA number distributions then generally the Markovian telegraph model is not a good approximation of the non-Markovian delay telegraph model.

In the main text, we used the delay telegraph model to infer the synthetic data from the synthetic data generated by the delay SSA. For comparison, we repeat the same but now we use the telegraph model (rather than the delay telegraph model) to calculate step (ii) in the inference algorithm, i.e. $P(k;\theta)$ in Eq. (4.7) in the main text is now chosen to be the steady-state solution of the telegraph model. Once the best parameter set $\theta^*$ is found, we calculate two scores: (i) the fitness given by the smallest negative log-likelihood value found by the optimizer normalized over the sample size. (ii) the mean relative error (for definition see Eq. (4.5) in the main text). In Fig. 3e, we show both of these scores obtained for 20 independent numerical experiments – clearly the error using the delay telegraph model for the inference algorithm is significantly lower than if the telegraph model is used.

These observations are further reinforced in Fig. 3f where we show the best fit distributions and the corresponding relative errors in the estimates of the burst size and the burst frequency (bar chart insets). These are computed using the formulae:

$$\begin{aligned} \alpha &= \text{Relative error in the burst frequency estimate} = \frac{|\sigma_{\text{on}}^* - \sigma_{\text{on}}|}{|\sigma_{\text{on}}^*|}, \\ \beta &= \text{Relative error in the burst size estimate} = \frac{|\rho^*/\sigma_{\text{off}}^* - \rho/\sigma_{\text{off}}|}{|\rho^*/\sigma_{\text{off}}^*|}. \end{aligned} \tag{2.3}$$

We note that the distributions are well fit in all cases, using both telegraph and delay telegraph models. However the errors $\alpha$ and $\beta$ are considerably larger for the former.

In Table 6 we show the true and estimated parameters for the 6 distributions in Fig. 3f. The estimates for $\rho$ are accurate independent of the choice of model; however the estimates for the promoter switching rates $\sigma_{\text{off}}, \sigma_{\text{on}}$ are far more accurate using the delay telegraph model. *Hence we conclude that although inference using the telegraph model provides a histogram that fits well with the synthetic data, the inferred parameter values have little meaning because they are not an accurate reflection of the true parameter values.*

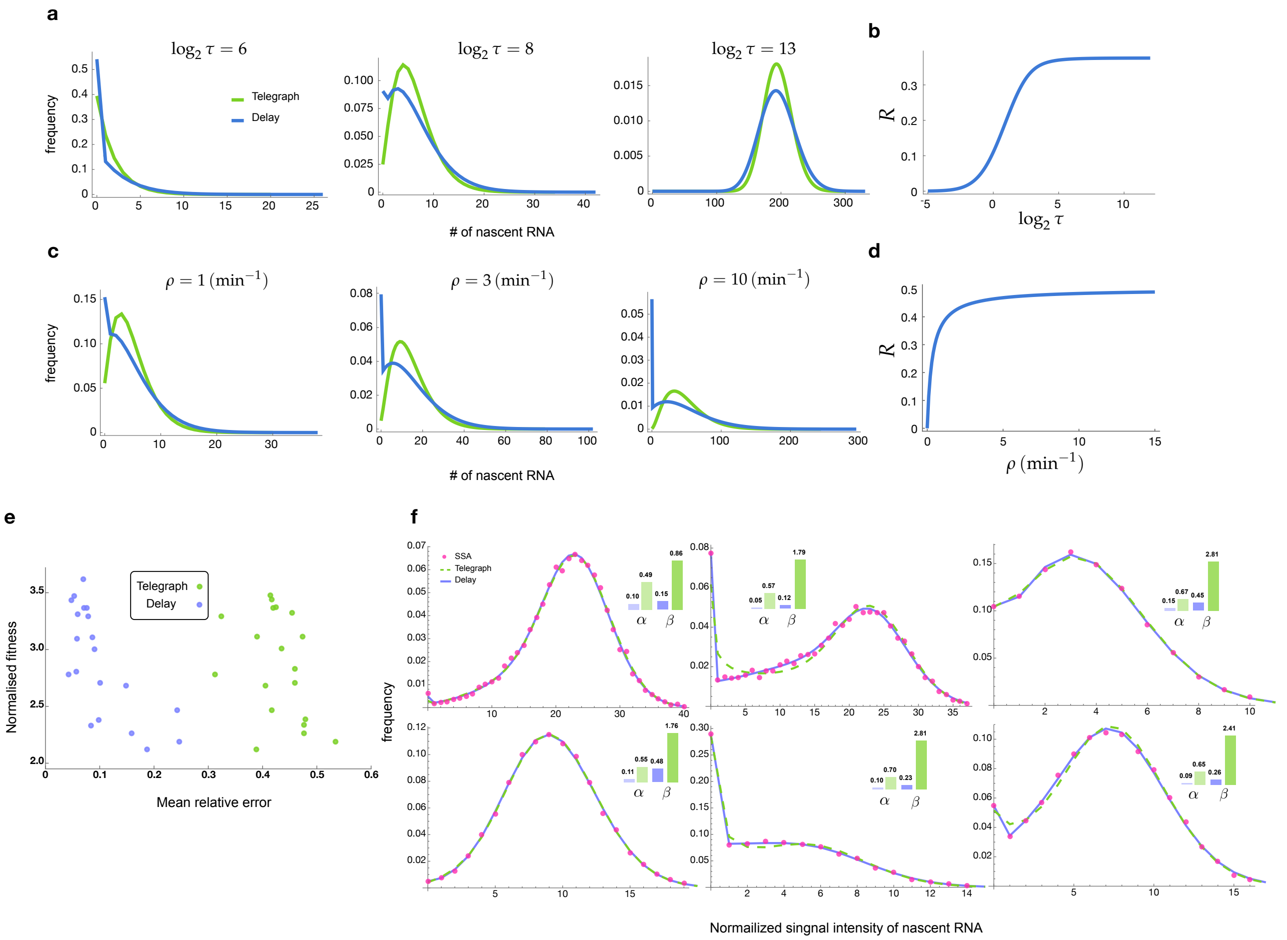

**Figure 3: a-d**. Comparison of the stochastic properties of the delay telegraph model and the telegraph model. **a.** Distributions of the nascent mRNA predicted by the delay telegraph model and the telegraph model for various values of $\tau$. We fix the parameters $(\sigma_{\text{off}}, \sigma_{\text{on}}, \rho) = (0.6, 0.03, 1)$ which implies that the change of $\tau$ leads to a change in $x = \tau(\sigma_{\text{on}} + \sigma_{\text{off}})$ at constant $y = (\rho/\sigma_{\text{off}})(\sigma_{\text{off}}/(\sigma_{\text{off}} + \sigma_{\text{on}}))^2$. **b.** Corresponding relative error $R$ between the variances of two models calculated as a function of $\tau$ using Eq. (2.2). **c.** Distributions of the nascent mRNA predicted by the delay telegraph model and the telegraph model for various values of $\rho$. We fix the parameters $(\sigma_{\text{off}}, \sigma_{\text{on}}, \tau) = (0.6, 0.03, 100)$ which implies that the change of $\rho$ leads to a change in $y$ at constant $x$. **d.** Corresponding relative error $R$ between the variances of two models calculated as a function of $\rho$ using Eq. (2.2). **e-f.** Inference of transcriptional parameters using as input synthetic fluorescent signal data generated by SSA simulations of transcription and fluorescent tagging for $10^4$ cells (see Methods section of the main text). **e.** Mean relative error and normalised fitness score (fitness/number of samples) plot for 20 sets of numerical experiments. The inference is done in two different ways, using either the telegraph model (green) or the delayed telegraph model (blue). **f.** Distributions of total fluorescent intensity from synthetic data (red dots) fit using the inference algorithm with telegraph model (dashed green) or delayed telegraph model (blue) for 6 different parameter sets. The insets show the relative errors in the estimates of the burst frequency ($\alpha$) and of the burst size ($\beta$) calculated using Eq. (2.3). Note that while both models provide a very good fit to the distribution from synthetic data, nevertheless parameter estimation is far more accurate using a delayed telegraph model. This is also reflected in (a) where we see low fitness scores for both models but a high mean relative error for estimates based on the telegraph model. The true and estimated parameters are shown in Table 6.

| | True | | | | | Delay | | | | | Telegraph | | | | |
|---|---|---|---|---|---|---|---|---|---|---|---|---|---|---|---|
| Parameter | $\sigma_{\text{off}}$ | $\sigma_{\text{on}}$ | $\rho$ | burst size | $f_{\text{ON}}$ | $\sigma_{\text{off}}$ | $\sigma_{\text{on}}$ | $\rho$ | burst size | $f_{\text{ON}}$ | $\sigma_{\text{off}}$ | $\sigma_{\text{on}}$ | $\rho$ | burst size | $f_{\text{ON}}$ |
| Set 1 | 1.05 | 8.20 | 57.99 | 55.09 | 0.89 | 0.94 | 7.19 | 57.92 | 61.87 | 0.88 | 0.60 | 4.10 | 59.51 | 99.42 | 0.87 |
| Set 2 | 1.27 | 3.14 | 58.17 | 45.69 | 0.71 | 1.13 | 2.91 | 58.01 | 51.16 | 0.72 | 0.45 | 1.22 | 59.43 | 133.07 | 0.73 |
| Set 3 | 2.27 | 5.80 | 12.55 | 5.53 | 0.72 | 1.60 | 4.48 | 12.21 | 7.63 | 0.74 | 0.59 | 1.71 | 12.09 | 20.61 | 0.74 |
| Set 4 | 1.18 | 9.94 | 24.92 | 21.15 | 0.89 | 0.94 | 8.74 | 24.74 | 26.37 | 0.90 | 0.46 | 4.12 | 24.92 | 54.00 | 0.90 |
| Set 5 | 2.26 | 2.14 | 18.64 | 8.26 | 0.49 | 1.77 | 1.76 | 18.28 | 10.36 | 0.50 | 0.54 | 0.58 | 17.60 | 32.59 | 0.52 |
| Set 6 | 1.38 | 4.77 | 21.74 | 15.79 | 0.78 | 1.08 | 4.00 | 21.50 | 19.94 | 0.79 | 0.38 | 1.49 | 21.31 | 56.44 | 0.80 |

**Table 6:** Estimates using the inference algorithm with delay telegraph and telegraph models for the six parameter sets in Fig. 3f.

### 3 Inference results using mature mRNA data

#### 3.1 Influence of segmentation on inference

In main text Section 2.2, we used independent segmentation tools to study segmentation artifacts. The inference results under segmentation 1, segmentation 2 and segmentation 2 without counting the transcriptional site are summarised in Table 7).

| | segmentation 1 | | | | | segmentation 2 | | | | | segmentation 2 without TS | | | | |
|---|---|---|---|---|---|---|---|---|---|---|---|---|---|---|---|
| Parameter | $\sigma_{\text{off}}$ | $\sigma_{\text{on}}$ | $\rho$ | burst size | $f_{\text{ON}}$ | $\sigma_{\text{off}}$ | $\sigma_{\text{on}}$ | $\rho$ | burst size | $f_{\text{ON}}$ | $\sigma_{\text{off}}$ | $\sigma_{\text{on}}$ | $\rho$ | burst size | $f_{\text{ON}}$ |
| Set 1 | 21.18 | 4.73 | 47.15 | 2.23 | 0.18 | 3.27 | 3.36 | 20.62 | 6.31 | 0.51 | 2.92 | 2.45 | 21.05 | 7.22 | 0.46 |
| Set 2 | 19.53 | 5.44 | 40.11 | 2.05 | 0.22 | 3.22 | 3.87 | 19.17 | 5.95 | 0.55 | 2.46 | 2.68 | 18.35 | 7.46 | 0.52 |
| Set 3 | 16.53 | 4.77 | 39.48 | 2.39 | 0.22 | 3.67 | 4.05 | 20.09 | 5.47 | 0.52 | 3.00 | 2.87 | 19.73 | 6.57 | 0.49 |
| Set 4 | 28.72 | 5.16 | 50.00 | 1.74 | 0.15 | 5.79 | 4.58 | 23.04 | 3.98 | 0.44 | 5.27 | 3.27 | 24.33 | 4.61 | 0.38 |
| Mean | 21.49 | 5.02 | 44.19 | 2.10 | 0.19 | 3.99 | 3.97 | 20.73 | 5.43 | 0.50 | 3.41 | 2.82 | 20.87 | 6.46 | 0.46 |
| Std | 5.19 | 0.34 | 5.21 | 0.28 | 0.03 | 1.06 | 0.43 | 1.43 | 0.89 | 0.04 | 1.26 | 0.35 | 2.56 | 1.29 | 0.06 |

**Table 7:** Inferred transcriptional parameters using merged mature mRNA data from segmentation 1, segmentation 2 and segmentation 2 without transcriptional site (TS).

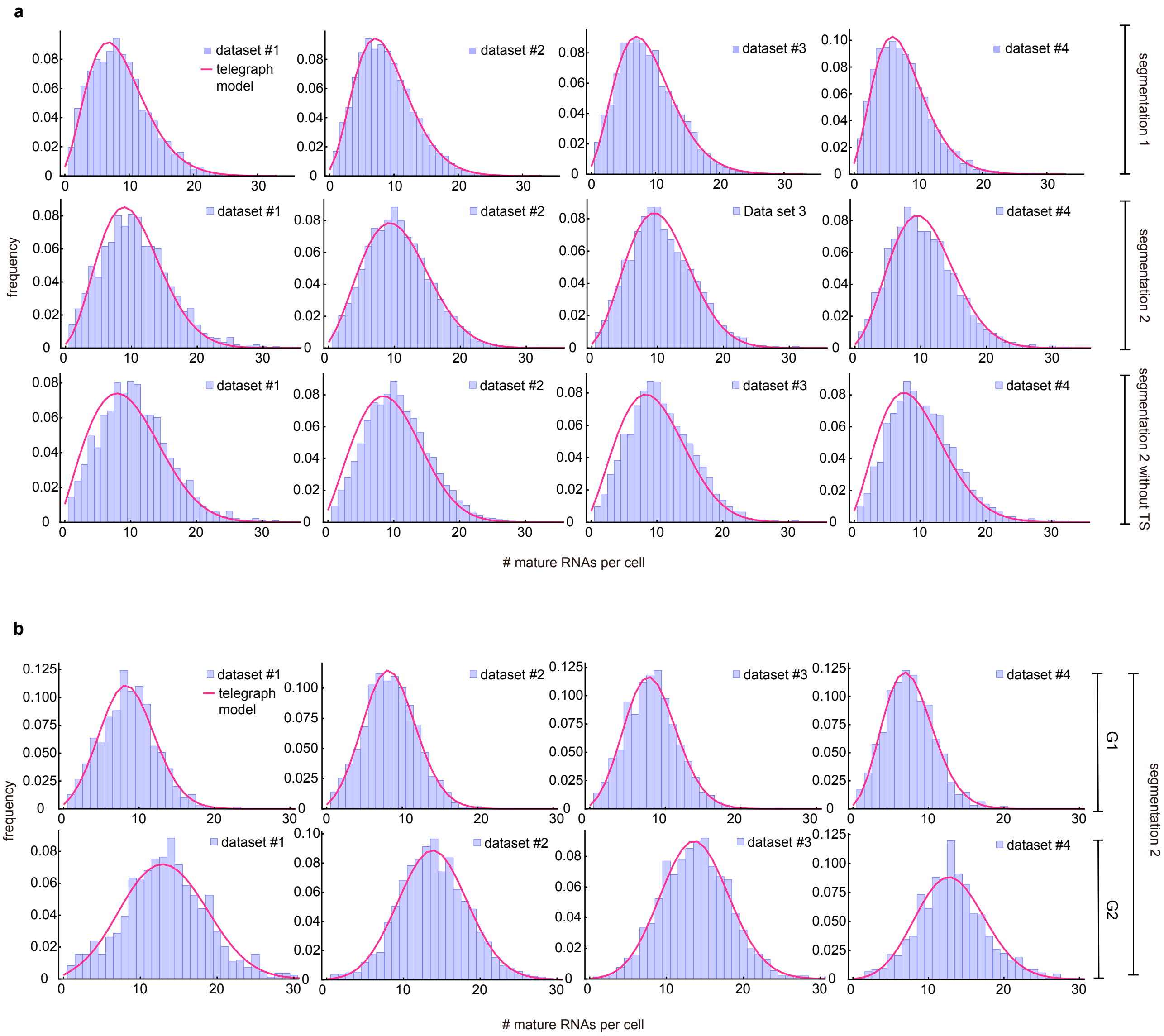

**Figure 4: a.** Merged mature mRNA count distribution (purple) under segmentation method 1 and with/without counting the transcriptional site (TS) under segmentation method 2 with a best fit obtained from the telegraph model (magenta curves). **b.** cell-cyle specific mature mRNA count distribution (purple) under segmentation method 2 with a best fit obtained from the telegraph model (magenta curves).

#### 3.2 Confidence intervals for estimates from merged and cell-cycle specific mRNA data using segmentation 2

We obtain 95% confidence intervals for the parameters estimated from mature mRNA data under segmentation 2 (shown in Fig. 3f of the main text). The confidence intervals are shown for $\sigma_{\text{off}}, \sigma_{\text{on}}, \rho$ in Table 8.

| mature | | $\sigma_{\text{off}}$ | $\sigma_{\text{on}}$ | $\rho$ | CI for $\sigma_{\text{off}}, \sigma_{\text{on}}, \rho$ | | |
|---|---|---|---|---|---|---|---|
| merged | Set 1 | 3.27 | 3.36 | 20.62 | (2.08, 6.08 ) | (2.81, 4.16 ) | (18.03, 25.97 ) |
| | Set 2 | 3.22 | 3.87 | 19.17 | (2.38, 4.69 ) | (3.41, 4.44 ) | (17.51, 21.62 ) |
| | Set 3 | 3.67 | 4.05 | 20.09 | (2.54, 5.92 ) | (3.51, 4.79 ) | (18.03, 23.67 ) |
| | Set 4 | 5.79 | 4.58 | 23.04 | (3.33, 13.33 ) | (3.79, 5.85 ) | (19.36, 33.51 ) |
| G1 | Set 1 | 0.69 | 2.76 | 10.71 | (0.27, 2.35 ) | (1.73, 4.96 ) | (9.77 , 12.85 ) |
| | Set 2 | 0.60 | 2.80 | 10.27 | (0.35, 1.24 ) | (2.13, 3.92 ) | (9.77 , 11.15 ) |
| | Set 3 | 0.88 | 3.41 | 10.33 | (0.38, 2.84 ) | (2.23, 5.77 ) | (9.58 , 12.35 ) |
| | Set 4 | 2.46 | 5.21 | 11.12 | (0.46, 300.00) | (2.68, 18.86) | (8.96 , 216.67) |
| G2 | Set 1 | 0.42 | 0.99 | 9.45 | (0.22, 0.91 ) | (0.65, 1.48 ) | (8.64 , 10.86 ) |
| | Set 2 | 0.14 | 1.01 | 7.92 | (0.05, 0.36 ) | (0.56, 1.72 ) | (7.61 , 8.46 ) |
| | Set 3 | 0.18 | 1.24 | 7.91 | (0.05, 0.72 ) | (0.57, 2.59 ) | (7.49 , 8.85 ) |
| | Set 4 | 0.42 | 1.68 | 8.19 | (0.12, 2.07 ) | (0.76, 3.50 ) | (7.41, 10.28) |

**Table 8:** 95% confidence interval intervals for the estimates from experimental mature mRNA data using segmentation 2

#### 3.3 Effect of synchronization of gene copies in G2 phase on parameter inference

In the main text, we assumed that the transcription of two allele copies in the G2 phase are independent. Here we consider the opposite scenario where the two allele copies in G2 phase are perfectly synchronized with each other, i.e. when one copy switches from on to off, the other copy also switches from on to off. Note that the time at which mRNA is transcribed from each copy is however not the same. Using this modified simulation algorithm, we re-perform the inference using the experimental data under segmentation 2. The results are shown in Table 9. Comparing these values with those estimated for phase G2 under the assumption of allele independence (main text Fig. 3f right panel), we find that $\rho$ and $f_{\text{ON}}$ are very similar but the other parameters vary considerably.

| mature | | $\sigma_{\text{off}}$ | $\sigma_{\text{on}}$ | $\rho$ | burst size | $f_{\text{ON}}$ |
|---|---|---|---|---|---|---|
| G2 sync | Set 1 | 0.62 | 2.17 | 8.45 | 13.69 | 0.78 |
| | Set 2 | 0.37 | 3.26 | 7.72 | 21.05 | 0.90 |
| | Set 3 | 0.67 | 4.42 | 7.92 | 11.85 | 0.87 |
| | Set 4 | 0.55 | 3.45 | 7.55 | 13.62 | 0.86 |
| | Mean | 0.55 | 3.32 | 7.91 | 15.05 | 0.85 |
| | Std | 0.13 | 0.92 | 0.39 | 4.09 | 0.05 |

**Table 9:** Inferred transcriptional rate (normalized) per gene copy for the G2 cell cycle phase under the assumption that the two gene states are perfectly synchronized.

### 4 Inference results using nascent mRNA data

#### 4.1 Inference using non-curated data

We show the inference results of the experimental non-curated nascent mRNA data (with and without taking into consideration the cell-cycle) for the four datasets (Table 10) and their 95% confidence interval using the profile likelihood estimate (Table 11). The best fit distributions for merged and cell-cycle specific data are shown in Fig. 5.

| nascent | | $\sigma_{\text{off}}$ | $\sigma_{\text{on}}$ | $\rho$ | burst size | $f_{\text{ON}}$ |
|---|---|---|---|---|---|---|
| merged | Set 1 | 5.24 | 5.07 | 77.46 | 14.78 | 0.49 |
| | Set 2 | 5.58 | 5.13 | 82.11 | 14.71 | 0.48 |
| | Set 3 | 5.11 | 5.17 | 76.09 | 14.90 | 0.50 |
| | Set 4 | 5.96 | 6.13 | 73.23 | 12.30 | 0.51 |
| | Mean | 5.47 | 5.37 | 77.22 | 14.17 | 0.50 |
| | Std | 0.38 | 0.50 | 3.70 | 1.25 | 0.01 |
| G1 | Set 1 | 1.11 | 3.76 | 37.83 | 34.10 | 0.77 |
| | Set 2 | 1.53 | 3.94 | 41.33 | 27.06 | 0.72 |
| | Set 3 | 0.95 | 3.23 | 36.79 | 38.56 | 0.77 |
| | Set 4 | 1.28 | 3.76 | 36.04 | 28.09 | 0.75 |
| | Mean | 1.22 | 3.67 | 38.00 | 31.95 | 0.75 |
| | Std | 0.25 | 0.31 | 2.34 | 5.39 | 0.02 |
| G2 | Set 1 | 0.74 | 1.69 | 35.00 | 47.30 | 0.70 |
| | Set 2 | 0.82 | 2.18 | 36.30 | 44.37 | 0.73 |
| | Set 3 | 0.91 | 2.19 | 34.54 | 37.90 | 0.71 |
| | Set 4 | 1.08 | 2.61 | 33.27 | 30.76 | 0.71 |
| | Mean | 0.89 | 2.17 | 34.78 | 40.08 | 0.71 |
| | Std | 0.15 | 0.38 | 1.25 | 7.35 | 0.01 |

**Table 10:** Estimated parameters from the non-curated distribution of the normalized intensity of the brightest nuclear spot (nascent mRNA data) constructed by merging all data or else specific to the cell cycle phases G1 and G2. The elongation time $\tau$ is estimated to be 0.785 mins, based on measurements of the elongation speed.

| nascent | | $\sigma_{\text{off}}$ | $\sigma_{\text{on}}$ | $\rho$ | burst size | $f_{\text{ON}}$ | CI for $\sigma_{\text{off}}, \sigma_{\text{on}}, \rho$ | | |
|---|---|---|---|---|---|---|---|---|---|
| merged | Set 1 | 5.24 | 5.07 | 77.46 | 14.78 | 0.49 | (4.42, 6.23) | (4.68, 5.45) | (73.48, 82.92) |
| | Set 2 | 5.58 | 5.13 | 82.11 | 14.71 | 0.48 | (5.05, 6.15) | (4.97, 5.37) | (79.38, 85.53) |
| | Set 3 | 5.11 | 5.17 | 76.09 | 14.90 | 0.50 | (4.53, 5.71) | (4.98, 5.38) | (73.49, 79.64) |
| | Set 4 | 5.96 | 6.13 | 73.23 | 12.30 | 0.51 | (5.19, 7.22) | (5.87, 6.57) | (69.34, 78.37) |
| G1 | Set 1 | 1.11 | 3.76 | 37.83 | 34.10 | 0.77 | (0.77, 1.66) | (3.04, 4.63) | (36.26, 39.96) |
| | Set 2 | 1.53 | 3.94 | 41.33 | 27.06 | 0.72 | (1.29, 1.88) | (3.64, 4.40) | (40.18, 42.85) |
| | Set 3 | 0.95 | 3.23 | 36.79 | 38.56 | 0.77 | (0.77, 1.17) | (2.86, 3.60) | (36.05, 37.69) |
| | Set 4 | 1.28 | 3.76 | 36.04 | 28.09 | 0.75 | (0.99, 1.73) | (3.18, 4.34) | (34.68, 37.75) |
| G2 | Set 1 | 0.74 | 1.69 | 35.00 | 47.30 | 0.70 | (0.54, 1.02) | (1.36, 2.08) | (33.64, 36.72) |
| | Set 2 | 0.82 | 2.18 | 36.30 | 44.37 | 0.73 | (0.66, 1.07) | (1.85, 2.52) | (35.35, 37.40) |
| | Set 3 | 0.91 | 2.19 | 34.54 | 37.90 | 0.71 | (0.70, 1.23) | (1.85, 2.61) | (33.38, 36.05) |
| | Set 4 | 1.08 | 2.61 | 33.27 | 30.76 | 0.71 | (0.75, 1.71) | (2.00, 3.41) | (31.91, 35.40) |

**Table 11:** 95% confidence intervals for non-curated data estimated using the profile likelihood method.

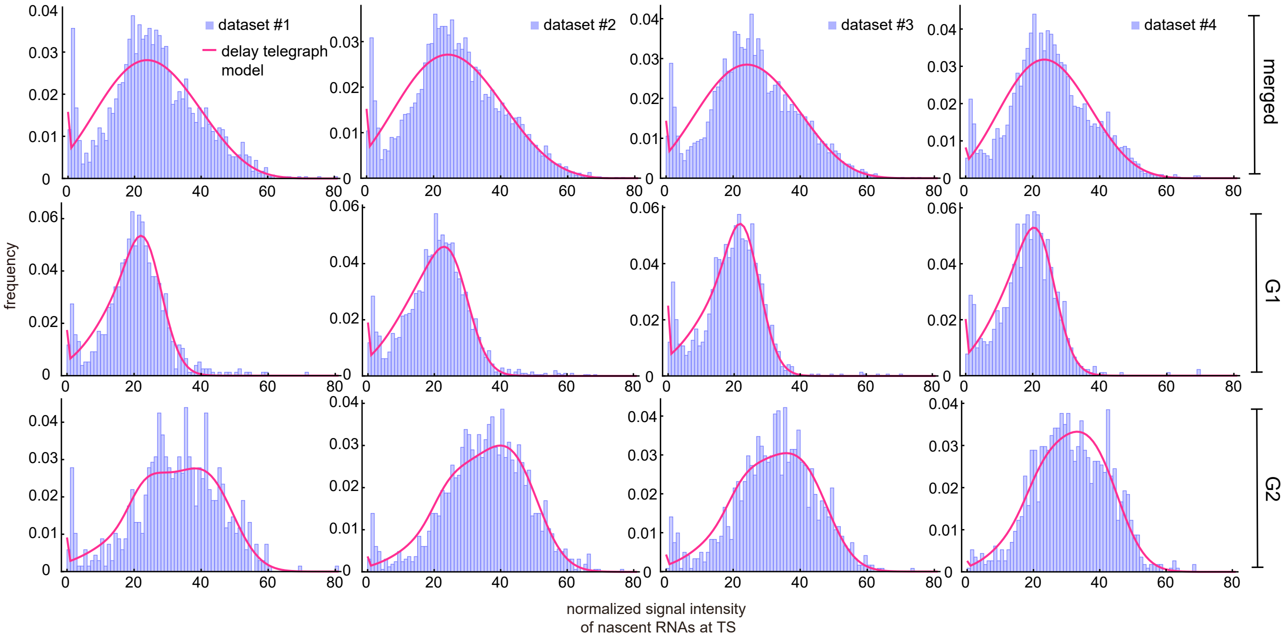

**Figure 5:** Inference results using merged and cell-cycle specific nascent data. Experimental distributions (purple) are fit using the delay telegraph model (magenta curves).

#### 4.2 Inference using curated data

We show the inference results using nascent mRNA data curated with the rejection method in Table 12 and Fig. 6, and with the fusion method in Table 13. We also used the profile likelihood method to obtain the 95% confidence intervals for the fusion method with $k = 4$; the results are shown in Table 14. We also reinferred the parameters in the G2 phase for fusion corrected data by assuming that the allele states are perfectly synchronised; these are shown in Table 15.

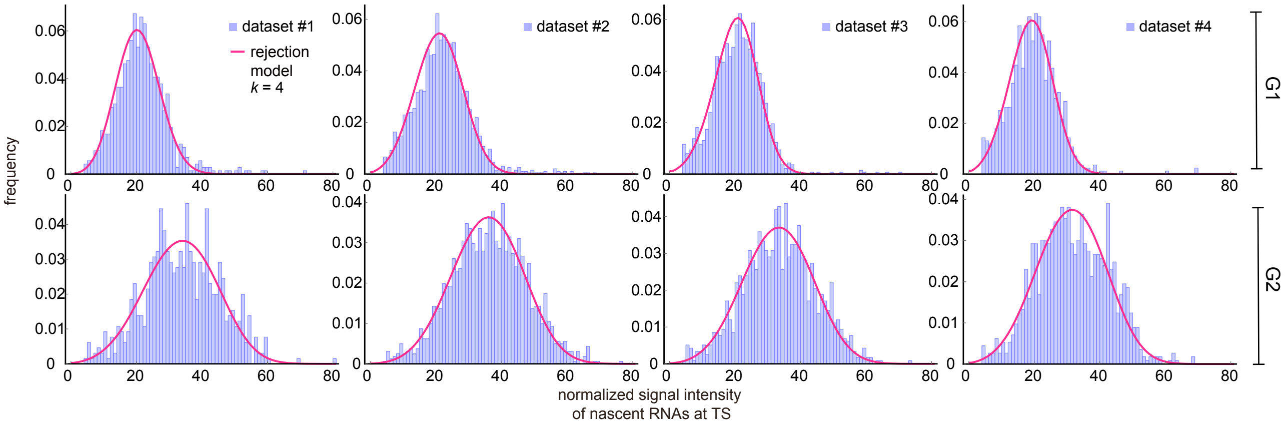

**Figure 6:** Inference results using cell-cycle specific data curated with the rejection method (only the distributions for $k = 4$ are shown). Corresponding fitted distributions (purple) for G1 (top row) and G2 (bottom row) using the delay telegraph model (magenta curves).

| $k=1$ | | $\sigma_{\text{off}}$ | $\sigma_{\text{on}}$ | $\rho$ | burst size | $f_{\text{ON}}$ |
|---|---|---|---|---|---|---|
| G1 | Set 1 | 1.06 | 4.33 | 36.62 | 34.63 | 0.80 |
| | Set 2 | 1.45 | 4.42 | 39.84 | 27.51 | 0.75 |
| | Set 3 | 1.04 | 3.84 | 36.66 | 35.19 | 0.79 |
| | Set 4 | 1.27 | 4.12 | 35.41 | 27.78 | 0.76 |
| | Mean | 1.21 | 4.18 | 37.13 | 31.28 | 0.78 |
| | Std | 0.19 | 0.26 | 1.90 | 4.20 | 0.02 |
| G2 | Set 1 | 0.84 | 2.10 | 34.54 | 41.20 | 0.71 |
| | Set 2 | 0.97 | 2.94 | 35.49 | 36.73 | 0.75 |
| | Set 3 | 0.94 | 2.53 | 33.68 | 35.99 | 0.73 |
| | Set 4 | 1.20 | 3.07 | 32.81 | 27.28 | 0.72 |
| | Mean | 0.99 | 2.66 | 34.13 | 35.30 | 0.73 |
| | Std | 0.15 | 0.44 | 1.15 | 5.82 | 0.02 |

| $k=3$ | | $\sigma_{\text{off}}$ | $\sigma_{\text{on}}$ | $\rho$ | burst size | $f_{\text{ON}}$ |
|---|---|---|---|---|---|---|
| G1 | Set 1 | 2.66 | 9.03 | 39.73 | 14.92 | 0.77 |
| | Set 2 | 2.55 | 7.41 | 42.04 | 16.46 | 0.74 |
| | Set 3 | 1.64 | 6.67 | 37.69 | 22.92 | 0.80 |
| | Set 4 | 2.22 | 7.32 | 37.02 | 16.65 | 0.77 |
| | Mean | 2.27 | 7.61 | 39.12 | 17.73 | 0.77 |
| | Std | 0.46 | 1.00 | 2.26 | 3.54 | 0.02 |
| G2 | Set 1 | 2.01 | 4.35 | 37.23 | 18.50 | 0.68 |
| | Set 2 | 2.16 | 5.10 | 38.56 | 17.85 | 0.70 |
| | Set 3 | 1.70 | 4.05 | 35.55 | 20.85 | 0.70 |
| | Set 4 | 1.88 | 4.16 | 34.48 | 18.30 | 0.69 |
| | Mean | 1.94 | 4.41 | 36.45 | 18.88 | 0.69 |
| | Std | 0.19 | 0.48 | 1.80 | 1.34 | 0.01 |

| $k=2$ | | $\sigma_{\text{off}}$ | $\sigma_{\text{on}}$ | $\rho$ | burst size | $f_{\text{ON}}$ |
|---|---|---|---|---|---|---|
| G1 | Set 1 | 1.58 | 6.33 | 37.77 | 23.89 | 0.80 |
| | Set 2 | 1.93 | 5.93 | 40.86 | 21.16 | 0.75 |
| | Set 3 | 1.28 | 5.15 | 37.07 | 29.07 | 0.80 |
| | Set 4 | 1.55 | 5.29 | 35.92 | 23.10 | 0.77 |
| | Mean | 1.59 | 5.67 | 37.91 | 24.30 | 0.78 |
| | Std | 0.27 | 0.55 | 2.11 | 3.38 | 0.02 |
| G2 | Set 1 | 1.43 | 3.42 | 35.83 | 25.11 | 0.71 |
| | Set 2 | 1.69 | 4.39 | 37.42 | 22.10 | 0.72 |
| | Set 3 | 1.33 | 3.38 | 34.64 | 26.08 | 0.72 |
| | Set 4 | 1.56 | 3.68 | 33.70 | 21.61 | 0.70 |
| | Mean | 1.50 | 3.72 | 35.39 | 23.72 | 0.71 |
| | Std | 0.16 | 0.47 | 1.61 | 2.20 | 0.01 |

| $k=4$ | | $\sigma_{\text{off}}$ | $\sigma_{\text{on}}$ | $\rho$ | burst size | $f_{\text{ON}}$ |
|---|---|---|---|---|---|---|
| G1 | Set 1 | 6.10 | 14.18 | 44.41 | 7.28 | 0.70 |
| | Set 2 | 3.72 | 9.57 | 43.96 | 11.81 | 0.72 |
| | Set 3 | 2.02 | 7.99 | 38.25 | 18.89 | 0.80 |
| | Set 4 | 2.75 | 8.60 | 37.78 | 13.76 | 0.76 |
| | Mean | 3.65 | 10.09 | 41.10 | 12.94 | 0.74 |
| | Std | 1.77 | 2.80 | 3.57 | 4.81 | 0.04 |
| G2 | Set 1 | 2.12 | 4.50 | 37.48 | 17.68 | 0.68 |
| | Set 2 | 2.59 | 5.68 | 39.55 | 15.25 | 0.69 |
| | Set 3 | 2.47 | 5.14 | 37.27 | 15.08 | 0.68 |
| | Set 4 | 2.18 | 4.55 | 35.17 | 16.13 | 0.68 |
| | Mean | 2.34 | 4.97 | 37.37 | 16.04 | 0.68 |
| | Std | 0.23 | 0.56 | 1.79 | 1.19 | 0.01 |

**Table 12:** (Rejection method) Estimated parameters by discarding the first $k$ signal bins of the experimental distribution of the signal intensity (and renormalizing afterwards). Inference is done for each of the four data sets. The elongation time is fixed to $\tau \approx 0.785$ mins.

| $k=1$ | | $\sigma_{\text{off}}$ | $\sigma_{\text{on}}$ | $\rho$ | burst size | $f_{\text{ON}}$ |
|---|---|---|---|---|---|---|
| G1 | Set 1 | 1.11 | 3.76 | 37.83 | 34.10 | 0.77 |
| | Set 2 | 1.53 | 3.94 | 41.33 | 27.06 | 0.72 |
| | Set 3 | 0.95 | 3.23 | 36.79 | 38.56 | 0.77 |
| | Set 4 | 1.28 | 3.76 | 36.04 | 28.09 | 0.75 |
| | Mean | 1.22 | 3.67 | 38.00 | 31.95 | 0.75 |
| | Std | 0.25 | 0.31 | 2.34 | 5.39 | 0.02 |
| G2 | Set 1 | 0.74 | 1.69 | 35.00 | 47.30 | 0.70 |
| | Set 2 | 0.82 | 2.18 | 36.30 | 44.37 | 0.73 |
| | Set 3 | 0.91 | 2.19 | 34.54 | 37.90 | 0.71 |
| | Set 4 | 1.08 | 2.61 | 33.27 | 30.76 | 0.71 |
| | Mean | 0.89 | 2.17 | 34.78 | 40.08 | 0.71 |
| | Std | 0.15 | 0.38 | 1.25 | 7.35 | 0.01 |

| $k=3$ | | $\sigma_{\text{off}}$ | $\sigma_{\text{on}}$ | $\rho$ | burst size | $f_{\text{ON}}$ |
|---|---|---|---|---|---|---|
| G1 | Set 1 | 0.71 | 2.79 | 36.38 | 51.39 | 0.80 |
| | Set 2 | 1.07 | 3.14 | 39.77 | 37.01 | 0.75 |
| | Set 3 | 0.63 | 2.35 | 35.74 | 56.84 | 0.79 |
| | Set 4 | 0.80 | 2.71 | 34.58 | 43.09 | 0.77 |
| | Mean | 0.80 | 2.75 | 36.62 | 47.08 | 0.78 |
| | Std | 0.19 | 0.32 | 2.23 | 8.78 | 0.02 |
| G2 | Set 1 | 0.52 | 1.25 | 34.30 | 65.86 | 0.71 |
| | Set 2 | 0.65 | 1.84 | 35.70 | 54.75 | 0.74 |
| | Set 3 | 0.71 | 1.81 | 33.85 | 47.67 | 0.72 |
| | Set 4 | 0.91 | 2.31 | 32.76 | 35.91 | 0.72 |
| | Mean | 0.70 | 1.80 | 34.15 | 51.05 | 0.72 |
| | Std | 0.16 | 0.43 | 1.22 | 12.57 | 0.01 |

| $k=2$ | | $\sigma_{\text{off}}$ | $\sigma_{\text{on}}$ | $\rho$ | burst size | $f_{\text{ON}}$ |
|---|---|---|---|---|---|---|
| G1 | Set 1 | 0.78 | 2.99 | 36.65 | 46.99 | 0.79 |
| | Set 2 | 1.14 | 3.27 | 40.01 | 35.06 | 0.74 |
| | Set 3 | 0.71 | 2.58 | 36.00 | 51.00 | 0.79 |
| | Set 4 | 0.96 | 3.09 | 35.08 | 36.52 | 0.76 |
| | Mean | 0.90 | 2.98 | 36.94 | 42.39 | 0.77 |
| | Std | 0.19 | 0.30 | 2.15 | 7.82 | 0.02 |
| G2 | Set 1 | 0.54 | 1.30 | 34.37 | 63.20 | 0.70 |
| | Set 2 | 0.67 | 1.87 | 35.74 | 53.70 | 0.74 |
| | Set 3 | 0.74 | 1.88 | 33.96 | 45.75 | 0.72 |
| | Set 4 | 0.94 | 2.36 | 32.84 | 34.89 | 0.72 |
| | Mean | 0.72 | 1.85 | 34.23 | 49.38 | 0.72 |
| | Std | 0.17 | 0.44 | 1.20 | 12.01 | 0.01 |

| $k=4$ | | $\sigma_{\text{off}}$ | $\sigma_{\text{on}}$ | $\rho$ | burst size | $f_{\text{ON}}$ |
|---|---|---|---|---|---|---|
| G1 | Set 1 | 0.69 | 2.73 | 36.30 | 52.85 | 0.80 |
| | Set 2 | 1.05 | 3.09 | 39.69 | 37.71 | 0.75 |
| | Set 3 | 0.63 | 2.35 | 35.74 | 56.78 | 0.79 |
| | Set 4 | 0.83 | 2.79 | 34.68 | 41.57 | 0.77 |
| | Mean | 0.80 | 2.74 | 36.60 | 47.23 | 0.78 |
| | Std | 0.19 | 0.31 | 2.17 | 9.04 | 0.02 |
| G2 | Set 1 | 0.56 | 1.34 | 34.45 | 61.05 | 0.70 |
| | Set 2 | 0.66 | 1.86 | 35.73 | 54.11 | 0.74 |
| | Set 3 | 0.67 | 1.72 | 33.70 | 50.56 | 0.72 |
| | Set 4 | 0.92 | 2.33 | 32.79 | 35.48 | 0.72 |
| | Mean | 0.70 | 1.81 | 34.17 | 50.30 | 0.72 |
| | Std | 0.15 | 0.41 | 1.24 | 10.80 | 0.01 |

**Table 13:** (Fusion method) Estimated parameters by combining the first $k$ signal bins of the experimental distribution of the signal intensity. Inference is done for each of the four data sets. The elongation time is fixed to $\tau \approx 0.785$ mins.

| nascent | | $\sigma_{\text{off}}$ | $\sigma_{\text{on}}$ | $\rho$ | burst size | $f_{\text{ON}}$ | CI for $\sigma_{\text{off}}, \sigma_{\text{on}}, \rho$ | | |
|---|---|---|---|---|---|---|---|---|---|
| G1 | Set 1 | 0.69 | 2.73 | 36.30 | 52.85 | 0.80 | (0.46, 1.05) | (2.12, 3.52) | (35.06, 37.91) |
| | Set 2 | 1.05 | 3.09 | 39.69 | 37.71 | 0.75 | (0.86, 1.29) | (2.73, 3.47) | (38.77, 40.92) |
| | Set 3 | 0.63 | 2.35 | 35.74 | 56.78 | 0.79 | (0.49, 0.79) | (1.99, 2.71) | (35.06, 36.58) |
| | Set 4 | 0.83 | 2.79 | 34.68 | 41.57 | 0.77 | (0.62, 1.16) | (2.27, 3.41) | (33.54, 36.01) |
| G2 | Set 1 | 0.56 | 1.34 | 34.45 | 61.05 | 0.70 | (0.40, 0.81) | (1.04, 1.73) | (33.35, 35.82) |
| | Set 2 | 0.66 | 1.86 | 35.73 | 54.11 | 0.74 | (0.51, 0.85) | (1.57, 2.21) | (34.87, 36.77) |
| | Set 3 | 0.67 | 1.72 | 33.70 | 50.56 | 0.72 | (0.50, 0.91) | (1.40, 2.11) | (32.78, 34.87) |
| | Set 4 | 0.92 | 2.33 | 32.79 | 35.48 | 0.72 | (0.62, 1.45) | (1.74, 3.09) | (31.65, 34.68) |

**Table 14:** Inference of the kinetic parameters $(\sigma_{\text{off}}, \sigma_{\text{on}}, \rho)$ using nascent data curated with the fusion method. Inferred values and the corresponding 95% confidence intervals of G1 and G2 cell-cycle specific data calculated using the profile likelihood method.

| nascent fusion | | $\sigma_{\text{off}}$ | $\sigma_{\text{on}}$ | $\rho$ | burst size | $f_{\text{ON}}$ |
|---|---|---|---|---|---|---|
| G2 sync | Set 1 | 1.84 | 4.42 | 33.84 | 18.42 | 0.71 |
| | Set 2 | 2.18 | 5.78 | 35.85 | 16.44 | 0.73 |
| | Set 3 | 2.15 | 5.36 | 33.61 | 15.63 | 0.71 |
| | Set 4 | 3.53 | 7.45 | 34.16 | 9.69 | 0.68 |
| | Mean | 2.42 | 5.75 | 34.36 | 15.05 | 0.71 |
| | Std. | 0.75 | 1.26 | 1.01 | 3.76 | 0.02 |

**Table 15:** Inferred transcriptional rate (normalized) per gene copy for the G2 cell cycle phase using nascent data curated with the fusion method under the assumption that the two gene states are perfectly synchronized.

### 5 Calculating the autocorrelation function (ACF) from stochastic simulations

Intensity traces of *GAL10* transcription were generated by a stochastic simulation algorithm (SSA) following the delay telegraph model presented in the main manuscript. Briefly, for each yeast cell, the ON and OFF *GAL10* promoter states are simulated such that they have an exponentially distributed lifetime with means $1/\sigma_{\text{off}}$ and $1/\sigma_{\text{off}}$, respectively. When the promoter was ON, the transcription of single *GAL10* mRNAs was initiated following a Poisson process with mean equal to the initiation rate $\rho$. The fluorescence trace $I(t)$, as measured experimentally over time, was obtained by convolving the train of mRNA initiation events with the trapezoidal intensity pulse of the PP7 tagging system (shown in Fig. 1c of the main text). It is important to note that this last step is necessary to accurately model the fluorescent intensities that are experimentally measured.

To account for cell cycle, we simulated an asynchronous population, where cells could switch from G1 to G2. In the latter, to account for two independent transcription sites, two independent intensity traces were generated and summed over time. The ratio of G1 to G2 cells was 1.15:1, 1.27:1, 1,24:1 and 0.80:1 for datasets 1, 2, 3 and 4 respectively, as determined from the smFISH experiments. It is important to note that: (i) live-cell experiments (as in [4]) are performed in asynchronous populations, without any sorting based on cell cycle phase; (ii) the average G1 phase duration for mother and daughter cells ($t_{\text{mother}} = 2400\,s$ and $t_{\text{daughter}} = 6480\,s$, respectively – unpublished data) is on the order of the typical total acquisition time (e.g. $1800\,s$). Hence, in our simulations, cells that were G1 at the start of the intensity trace (i.e. live-cell experiment) were allowed to switch to G2 before the end of the experiment. Trivially, cells that were G2 at the start remained so until the end of the experiment.

The autocorrelation function (ACF) was calculated as:

$$G(\Delta t) = \frac{\langle \delta I(t) \delta I(t + \Delta t) \rangle}{\langle I(t) \rangle^2} - 1, \tag{5.1}$$

where $\Delta t$ is the time-lag (e.g. $30\,s$ in [4]), $\langle \cdot \rangle$ denotes the temporal average over $t$, and $\delta I(t) = I(t) - \langle I(t) \rangle$. Fig. 7 shows the ACFs from Fig. 6c of the main manuscript, in which a linear fit is performed to correct for non-stationary effects, i.e. switching from G1 to G2 during the experiment, bleaching, heterogeneous galactose induction.

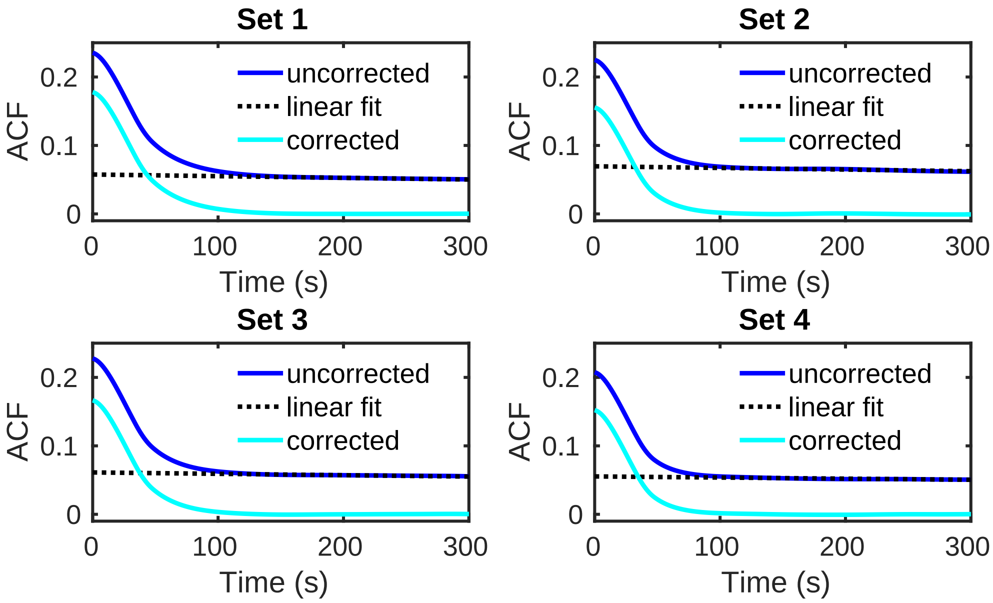

**Figure 7:** Autocorrelation functions of $10^4$ simulated *GAL10* intensity traces (solid blue lines). The transcriptional parameters for G1 and G2 cells in the 4 sets of experimental data were obtained using the fusion method (see Fig. 6d of the main manuscript). A linear fit (dashed black line) was subtracted to correct the ACFs for non-stationary switching from G1 to G2 (solid cyan lines).

### 6 Derivation of the exact solution for the master equation of the delay telegraph model and finite state projection

#### 6.1 Exact solution

The delay telegraph model includes four reactions

$$G \xrightarrow{\rho} G + N, \quad G \underset{\sigma_{\text{on}}}{\overset{\sigma_{\text{off}}}{\rightleftharpoons}} G^{\star}, \quad N \underset{\tau}{\Rightarrow} \varnothing, \tag{6.1}$$

where $G$ and $G^{\star}$ stand for the active (ON) and inactive (OFF) gene state, respectively, and $\sigma_{\text{off}}$ and $\sigma_{\text{on}}$ are the activation and inactivation rates, respectively. Once a nascent mRNA is produced, it will be removed from the system after a deterministic time $\tau$. We aim to find the quantitative description of the kinetics of nascent mRNA ($N$). We proceed by deriving the delay chemical master equation (CME) of Eq. (6.1). The same delay models have also been studied by other authors [5].

Let $P(0, n, t)$ and $P(1, n, t)$ be the probability of observing $n$ nascent mRNAs while the gene state is inactive and active at time $t$, respectively. Consequently, we can state

$$\begin{aligned} P(i, n, t + \Delta t) = & \{\text{Part A: Probability of one instant reaction occurring during } \Delta t\} \\ & + \{\text{Part B: Probability of one delayed reaction occurring during } \Delta t\} \\ & + \{\text{Part C: Probability of no reactions occurring during } \Delta t\}. \end{aligned} \tag{6.2}$$

Note that by instant reactions, we mean all reactions except the delayed reaction that removes nascent mRNA. Since Part A includes all Markovian reactions, it follows from standard arguments [6] that

$$\begin{aligned} P(0, n, t + \Delta t; 1, n, t) &= \sigma_{\text{off}} \Delta t P(1, n, t) + o(\Delta t), \\ P(1, n, t + \Delta t; 0, n, t) &= \sigma_{\text{on}} \Delta t P(0, n, t) + o(\Delta t), \\ P(1, n, t + \Delta t; 1, n-1, t) &= \rho \Delta t P(1, n-1, t) + o(\Delta t). \end{aligned} \tag{6.3}$$

The contribution of Part B requires a careful consideration of the history of the process: (i) $(1, n', t-\tau)$ leads to $(1, n'+1, t-\tau+\Delta t)$ when the gene was ON; (ii) $(1, n'+1, t-\tau+\Delta t)$ leads to $(i, n+1, t)$ where the gene state $i$ can be either OFF or ON; (iii) $(i, n+1, t)$ leads to $(i, n, t)$. Indeed, since the removal of nascent mRNA is a delayed reaction, (iii) is due to a production reaction in (i) a time interval $\tau$ earlier. The probability of (i) occurring is $\rho \Delta t P(1, n', t-\tau)$. The probability of (iii) occurring is 1 since every production event is followed by a removal event a time $\tau$ later. The probability of (ii) occurring is:

$$P(i, n\mathbf{+1}, t | 1, n'\mathbf{+1}, t - \tau + \Delta t) \tag{6.4}$$

As for Eq. (6.4), the two "**+1**"s means that the new nascent mRNA was produced during the time interval $(t-\tau, t-\tau+\Delta t)$, and it did not participate in any other reactions, thereby not influencing the probability at time $t + \Delta t$. In short,

$$P(i, n\mathbf{+1}, t | 1, n'\mathbf{+1}, t - \tau + \Delta t) = P(i, n, t | 1, n', t - \tau + \Delta t). \tag{6.5}$$

Furthermore, all $n'$ nascent mRNAs must leave the system prior to time $t$ as they were all born before time $t - \tau$. This implies that

$$P(i, n, t | 1, n', t - \tau + \Delta t) = P(i, n, t | 1, 0, t - \tau + \Delta t). \tag{6.6}$$

Hence the probability that event B occurs is given by the product of the probability of events (i), (ii) and (iii) and summing over all values of $n'$

$$\begin{aligned} P(i, n, t + \Delta t; i, n+1, t) &= \rho \Delta t \sum_{n'} P(1, n', t - \tau) P(i, n, t | 1, 0, t - \tau + \Delta t) \\ &= \rho \Delta t P(1, t - \tau) P(i, n, t | 1, 0, t - \tau + \Delta t), \end{aligned} \tag{6.7}$$

where we used $\sum_{n'} P(1, n', t - \tau) = P(1, t - \tau)$.

Finally, the contribution of Part C is obtained from simple probability arguments

$$\begin{aligned} P(0, n, t; 0, n, t) &= P(0, n, t) - P(1, n, t; 0, n, t) - P(0, n-1, t; 0, n, t) \\ &= P(0, n, t) - \sigma_{\text{on}} \Delta t P(0, n, t) - \rho \Delta t P(1, t - \tau) P(0, n-1, t | 1, 0, t - \tau + \Delta t), \\ P(1, n, t; 1, n, t) &= P(1, n, t) - P(0, n, t; 1, n, t) - P(1, n+1, t; 1, n, t) - P(1, n-1, t; 1, n, t) \\ &= P(1, n, t) - \sigma_{\text{off}} \Delta t P(1, n, t) - \rho \Delta t P(1, n, t) - \rho \Delta t P(1, t - \tau) P(1, n-1, t | 1, 0, t - \tau + \Delta t), \end{aligned} \tag{6.8}$$

where we used the same argument as in Eq. (6.7). Using Eqs. (6.2), (6.3), (6.7) and (6.8) and taking the limit of small $\Delta t$, we finally obtain the set of delay CMEs

$$\begin{cases} \dfrac{\mathrm{d}P(0,n,t)}{\mathrm{d}t} = -\sigma_{\text{on}}P(0,n,t) + \sigma_{\text{off}}P(1,n,t) \\ \qquad\qquad + (E-1)P(0,n-1,t|1,0,t-\tau)\rho P(1,t-\tau), \\ \dfrac{\mathrm{d}P(1,n,t)}{\mathrm{d}t} = \sigma_{\text{on}}P(0,n,t) - \sigma_{\text{off}}P(1,n,t) + \rho(E^{-1}-1)P(1,n,t) \\ \qquad\qquad + (E-1)P(1,n-1,t|1,0,t-\tau)\rho P(1,t-\tau), \end{cases} \tag{6.9}$$

where $E^k P(i,n,t) = P(i,n+k,t)$ is the step operator. Summing over all possible $n$ for the two equations in Eq. (6.9), we obtain

$$\begin{cases} \dfrac{\mathrm{d}P(0,t)}{\mathrm{d}t} = -\sigma_{\text{on}}P(0,t) + \sigma_{\text{off}}P(1,t), \\ \dfrac{\mathrm{d}P(1,t)}{\mathrm{d}t} = \sigma_{\text{on}}P(0,t) - \sigma_{\text{off}}P(1,t), \end{cases}$$

initiated with the inactive state, in which $P(0,t)$ and $P(1,t)$ are the probabilities of finding a cell at the inactivated and activated state at time $t$, and their solutions are

$$P(0,t) = \frac{\sigma_{\text{off}}}{\sigma_{\text{off}}+\sigma_{\text{on}}} + \frac{\sigma_{\text{on}}}{\sigma_{\text{off}}+\sigma_{\text{on}}} e^{-(\sigma_{\text{on}}+\sigma_{\text{off}})t}, \quad P(1,t) = \frac{\sigma_{\text{on}}}{\sigma_{\text{off}}+\sigma_{\text{on}}}\left(1 - e^{-(\sigma_{\text{on}}+\sigma_{\text{off}})t}\right).$$

Besides, it is noted that in Eq. (6.9)

$$P(1,t-\tau) = \frac{\sigma_{\text{on}}}{\sigma_{\text{off}}+\sigma_{\text{on}}}\left(1 - e^{-(\sigma_{\text{on}}+\sigma_{\text{off}})(t-\tau)}\right) =: h(t-\tau), \tag{6.10}$$

for $t \geq \tau$, and $h(t-\tau) = 0$ if $t < \tau$. The $n$ nascent mRNAs of the two conditional probabilities $P(i,n,t|1,0,t-\tau)$ for $i = 0,1$ in Eq. (6.9) are produced during time $(t-\tau,t)$, and the pertinent dynamics are that of a reaction system only composed of the three instant reactions in Eq. (6.1). Specifically, we have

$$\begin{cases} \dfrac{\mathrm{d}\widetilde{P}(0,n,t)}{\mathrm{d}t} = -\sigma_{\text{on}}\widetilde{P}(0,n,t) + \sigma_{\text{off}}\widetilde{P}(1,n,t), \\ \dfrac{\mathrm{d}\widetilde{P}(1,n,t)}{\mathrm{d}t} = \sigma_{\text{on}}\widetilde{P}(0,n,t) - \sigma_{\text{off}}\widetilde{P}(1,n,t) + \rho(E^{-1}-1)\widetilde{P}(1,n,t), \end{cases} \tag{6.11}$$

where the initial values are $\widetilde{P}(0,n,0) = 0$ for any $n$, $\widetilde{P}(1,n,0) = 1$ for $n = 0$ and equal to 0 otherwise, as well as $P(i,n,t|1,0,t-\tau) = \widetilde{P}(i,n,\tau)$ for any $i = 0,1$.

Let $G_i(u,t) = \sum_n (u+1)^n P(i,n,t)$ and $\widetilde{G}_i(u,t) = \sum_n (u+1)^n \widetilde{P}(i,n,t)$. We particularly define the generating function in such a form to simplify the notation. Then, using Eqs. (6.9), (6.10), and (6.11) we obtain

$$\begin{cases} \partial_t G_0 = -\sigma_{\text{on}}G_0 + \sigma_{\text{off}}G_1 - \rho h(t-\tau)u\mathbb{1}_{[\tau,\infty)}\widetilde{G}_0^{\tau}, \\ \partial_t G_1 = \rho u G_1 + \sigma_{\text{on}}G_0 - \sigma_{\text{off}}G_1 - \rho h(t-\tau)u\mathbb{1}_{[\tau,\infty)}\widetilde{G}_1^{\tau}, \end{cases} \tag{6.12}$$

and

$$\begin{cases} \partial_t \widetilde{G}_0 = -\sigma_{\text{on}}\widetilde{G}_0 + \sigma_{\text{off}}\widetilde{G}_1, \\ \partial_t \widetilde{G}_1 = \rho u \widetilde{G}_1 + \sigma_{\text{on}}\widetilde{G}_0 - \sigma_{\text{off}}\widetilde{G}_1, \end{cases} \tag{6.13}$$

where the arguments $u$ and $t$ in the generating functions are suppressed for clarity, and the superscript $\tau$ is used to emphasize the generating function $\widetilde{G}_i$ up to a particular time $\tau$. The initiation condition of Eq. (6.13) is $\widetilde{G}_0 = 0$ and $\widetilde{G}_1 = 1$ when $t = 0$.

Under the condition of steady-state as $t \to \infty$, Eq. (6.12) reduces to

$$\begin{cases} 0 = -\sigma_{\text{on}}G_0 + \sigma_{\text{off}}G_1 - \rho\bar{h}u\widetilde{G}_0^{\tau}, \\ 0 = \rho u G_1 + \sigma_{\text{on}}G_0 - \sigma_{\text{off}}G_1 - \rho\bar{h}u\widetilde{G}_1^{\tau}, \end{cases} \tag{6.14}$$

with $\bar{h} = \sigma_{\text{on}}/(\sigma_{\text{on}}+\sigma_{\text{off}})$.

Therefore, by solving Eq. (6.14) we obtain

$$\boxed{\begin{aligned} &G(u,\infty) = G_0(u,\infty) + G_1(u,\infty) \\ &= \frac{e^{-\frac{1}{2}\tau\left(\sqrt{\Delta(u)}+\sigma_{\text{off}}+\sigma_{\text{on}}-u\rho\right)}}{2\sqrt{\Delta(u)}\left(\sigma_{\text{off}}+\sigma_{\text{on}}\right)} \\ &\times \left[ \left(\sigma_{\text{off}}+\sigma_{\text{on}}\right)\left(\sqrt{\Delta(u)} + e^{\sqrt{\Delta(u)}\tau}\left(\sqrt{\Delta(u)}+\sigma_{\text{off}}+\sigma_{\text{on}}\right) - \sigma_{\text{off}} - \sigma_{\text{on}}\right) - u\left(e^{\sqrt{\Delta(u)}\tau} - 1\right)\rho\left(\sigma_{\text{off}} - \sigma_{\text{on}}\right)\right], \end{aligned}} \tag{6.15}$$

where we define $\Delta(u) = (\sigma_{\text{off}} - u\rho)^2 + 2\sigma_{\text{on}}(\sigma_{\text{off}} + u\rho) + \sigma_{\text{on}}^2$. Finally, we obtain the probability distribution by the following relation

$$P(n,t) = \frac{1}{n!}\partial_u^n G(u,\infty)|_{u=-1}, \quad \text{for all } n \in \mathbb{N}.$$

We checked that the obtained distribution agrees exactly with the nascent mRNA distribution of Eq. 7 in [3].

#### 6.2 Finite State Projection

The finite state projection (FSP) method [7] is an efficient numerical method that can be implemented to solve the CME (6.9) to any desired degree of accuracy. Because Eq. (6.9) is coupled with Eq. (6.11), here we show how to use a two-step FSP method to obtain the probability distribution.

By assuming that there exists a large $N$ such that $P(i,n,t) = 0$ and $\tilde{P}(i,n,t) = 0$ for any $i = 0,1$ and $n \geq N+1$, we denote

$$\mathbf{P}(t) = (P(0,0,t), P(0,1,t), \ldots, P(0,N,t), P(1,0,t), P(1,1,t), \ldots, P(1,N,t))^T$$

as a $N^2$-dimensional vector and $\tilde{\mathbf{P}}(t) \in \mathbb{R}^{N^2}$ similarly. We first use FSP method to solve Eq. (6.11) with the following ordinary differential equation (ODE) up to time $\tau$

$$\frac{\mathrm{d}}{\mathrm{d}t}\tilde{\mathbf{P}}(t) = \mathbf{A}\tilde{\mathbf{P}}(t) \tag{6.16}$$

where $\mathbf{A}$ is defined as

$$\mathbf{A} = \begin{pmatrix} -\sigma_{\text{on}}\mathbf{I}_N & 0 \\ 0 & \sigma_{\text{off}}\mathbf{I}_N + \mathbf{B} \end{pmatrix}_{2(N+1)\times 2(N+1)}, \quad \mathbf{B} = \begin{pmatrix} -\rho & 0 & \cdots & 0 & 0 \\ \rho & -\rho & \cdots & 0 & 0 \\ \vdots & \vdots & \ddots & \vdots & \vdots \\ 0 & 0 & \cdots & \rho & -\rho \end{pmatrix}_{N+1\times N+1}$$

with the initial condition

$$\tilde{\mathbf{P}}(0) = (\underbrace{0,0,\ldots,0}_{N+1},\underbrace{1,0,\ldots,0}_{N+1})^T.$$

Secondly, we solve the following inhomogeneous ODE

$$\frac{\mathrm{d}}{\mathrm{d}t}\mathbf{P}(t) = \mathbf{A}\mathbf{P}(t) + h(t-\tau)\left(\tilde{\mathbf{P}}(\tau) - \mathcal{S}\tilde{\mathbf{P}}(\tau)\right), \tag{6.17}$$

where $\mathcal{S}$ is the right-shift operator, i.e. for any $\mathbf{x} = (x_1, x_2, \ldots, x_N)$, $\mathcal{S}\mathbf{x} = (0, x_1, x_2, \ldots, x_{N-1})$ and

$$h(t) = \begin{cases} \frac{\rho\sigma_{\text{on}}}{\sigma_{\text{off}}+\sigma_{\text{on}}}\left(1 - e^{-(\sigma_{\text{on}}+\sigma_{\text{off}})t}\right) & t \geq 0, \\ 0 & t < 0, \end{cases}$$

with the initial condition

$$\mathbf{P}(0) = (\underbrace{1,0,\ldots,0}_{N+1},\underbrace{0,0,\ldots,0}_{N+1})^T.$$

Hence, the solution of Eq. (6.17) gives the approximate probability distribution $\mathbf{P}(t)$ for any $t > 0$.

Note that if one uses the generating function Eq. (6.15) to compute the probability distribution, the computation for the high-order derivatives can cause numerical instabilities due to lack of arithmetic precision – compare the exact solutions with precision 85 and 300 in the left and right panels of Fig. 8. Because FSP avoids the computation of the higher-order derivatives, it can be as numerically stable as computing the exact solution with high precision (right panel of Fig. 8). In Table 16, we show the computational efficiency of FSP versus numerical evaluation of the exact solution – FSP and the exact solution with high precision are hence comparable both in terms of accuracy and efficiency. In the main text, we use FSP.

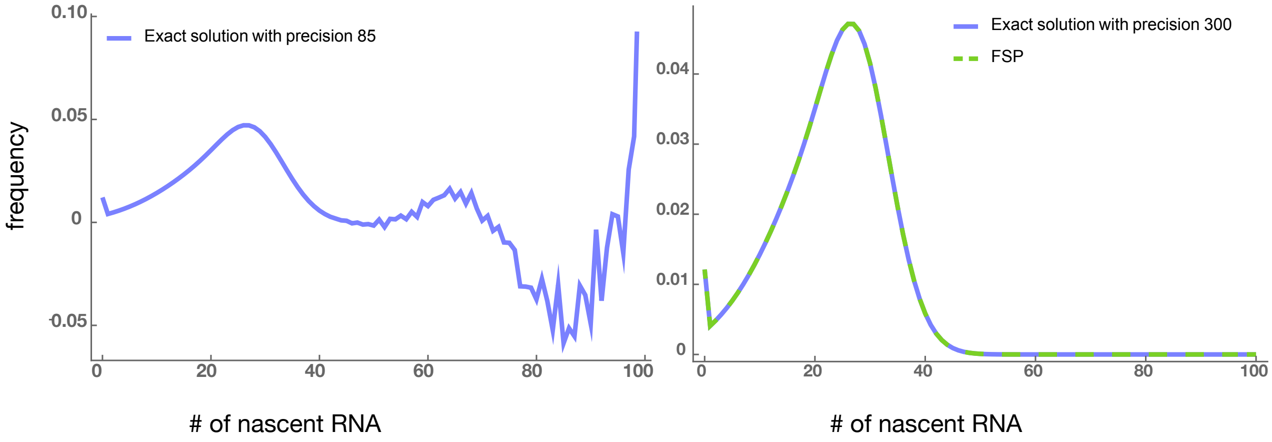

**Figure 8:** Left panel: Numerical instabilities due to the calculation of higher-order derivatives in the exact solution appear when the arithmetic precision is not very high (85). Right panel: These instabilities disappear when the precision increases to 300. The exact solution with such high precision agrees well with the FSP method using double-precision floating-point (Float64) type. The parameters are $\sigma_{\text{off}} = 1.71$, $\sigma_{\text{on}} = 5.82$, $\rho = 53.74$, and $\tau = 0.56$.

| | Exact solution: precision = 85 | Exact solution: precision = 300 | FSP method |
|---|---|---|---|
| Minimum time: | 6.422 ms | 9.277 ms | 8.317 ms |
| Median time: | 6.868 ms | 9.562 ms | 9.092 ms |
| Mean time: | 8.279 ms | 11.482 ms | 9.415 ms |
| Maximum time: | 16.791 ms | 16.919 ms | 14.203 ms |
| # of Simulations: | 604 | 436 | 531 |

**Table 16:** Comparison of the performance of three methods to compute the probability distribution. Parameters same as in Fig. 8. The time was calculated using the *Julia* package *BenchmarkTools.jl*.

### 7 Classification of the cell cycle phase: bimodal Gaussian distribution method versus the Fried/Baisch model

The distribution of DNA content intensities (DAPI) was fit using a bimodal distribution; those cells whose intensity was around one peak were classified as G1 and those around the second peak were classified as G2 (main text Fig 3e). We also did the classification using an alternative method, based on the Freid/Baisch model, which was recently employed in [8]. In Fig. 9 we show the distributions of fluorescent signal intensity for cells in the G1 phase (top row) and for those in the G2 phase (bottom row) for the four data sets described in the main text. Note that the method (bimodal Gaussian or Freid/Baisch) used to classify the cells in G1/G2 does not alter the intensity distribution hence verifying the robustness of our classification method.

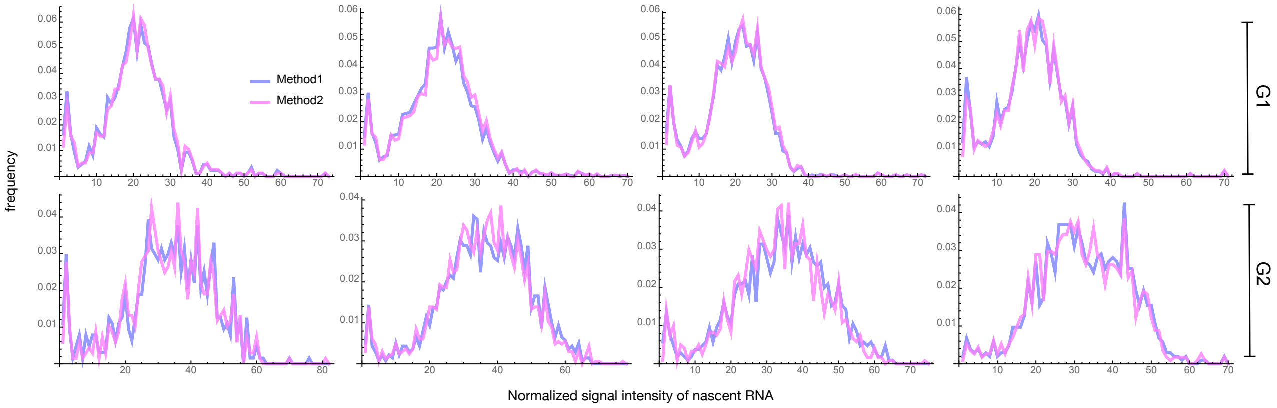

**Figure 9:** Blue curves (Method 1) show the fluorescent intensity distributions of the four experimental data sets after the classification of cells into G1 and G2 phases using the Fried/Baisch model. Magenta curves (Method 2) show the same but using a bimodal Gaussian, as described in the main text.

### References

[1] Kreutz, C., Raue, A., Kaschek, D. & Timmer, J. Profile likelihood in systems biology. *The FEBS journal* **280**, 2564–2571 (2013).

[2] Peccoud, J. & Ycart, B. Markovian modeling of gene-product synthesis. *Theoretical Population Biology* **48**, 222–234 (1995).

[3] Xu, H., Skinner, S. O., Sokac, A. M. & Golding, I. Stochastic Kinetics of Nascent RNA. *Physical Review Letters* **117**, 1–6 (2016).

[4] Donovan, B. T. *et al.* Live-cell imaging reveals the interplay between transcription factors, nucleosomes, and bursting. *The EMBO journal* **38**, e100809 (2019).

[5] Xu, H., Skinner, S. O., Sokac, A. M. & Golding, I. Stochastic kinetics of nascent rna. *Physical review letters* **117**, 128101 (2016).

[6] Gillespie, D. T. Stochastic simulation of chemical kinetics. *Annu. Rev. Phys. Chem.* **58**, 35–55 (2007).

[7] Munsky, B. & Khammash, M. The finite state projection algorithm for the solution of the chemical master equation. *J Chem Phys* **124**, 044104 (2006).

[8] Skinner, S. O. *et al.* Single-cell analysis of transcription kinetics across the cell cycle. *eLife* **5**, 1–24 (2016).
